# Mechanical Tension in Syndecan-1 is Regulated by Extracellular Mechanical Cues and Fluidic Shear Stress

**DOI:** 10.1101/2020.05.09.085894

**Authors:** Victoria Le, Lei Mei, Peter L. Voyvodic, Chi Zhao, David J. Busch, Jeanne C. Stachowiak, Aaron B. Baker

## Abstract

The endothelium plays a central role in regulating vascular homeostasis and is key in determining the response to materials implanted in the vascular system. Endothelial cells are uniquely sensitive to biophysical cues from applied forces and their local cellular microenvironment. The glycocalyx is a layer of proteoglycans, glycoproteins and glycosaminoglycans that lines the luminal surface of the vascular endothelium, interacting directly with the components of the blood and the forces of blood flow. In this work, we examined the changes in mechanical tension of syndecan-1, a cell surface proteoglycan that is an integral part of the glycocalyx, in response to substrate stiffness and fluidic shear stress. Our studies demonstrate that syndecan-1 is mechanically responsive to extracellular mechanical cues and alters its association with cytoskeletal and adhesion-related proteins in response to substrate stiffness and physiological flow.

## Introduction

The cells of the vascular system are subjected to complex biophysical cues that regulate homeostasis and pathophysiological processes^1,2^. These mechanical cues are sensed by many mechanisms including cell surface receptors, stretch sensitive ion channels and cytoskeleton-linked proteins^3^. In therapeutic applications, a key challenge is the development of engineered materials that can potentiate the curative effects of endogenous or delivered cells to enable the maximal therapeutic benefit from the implanted cells and material. It is increasingly recognized that mechanical cues including substrate rigidity and the presence of nanotopological features in the cell microenvironment can profoundly alter cell behavior and differentiation^4–8^.

The endothelial glycocalyx is a layer of proteoglycans, glycoproteins and glycans that lines the lumen of the vessels of the vascular system. This complex structure interacts directly with the forces of blood flow and serves as a key regulator of vascular permeability and interactions with circulating immune cells^9–11^. One component of the glycocalyx is syndecan-1 (SDC1), a cell surface proteoglycan that is post-translationally modified with heparan sulfate and chondroitin sulfate polysaccharide chains. Our group has recently identified SDC1 as a mechanosensor of shear stress in endothelial cells^11^, a mediator in endothelial inflammation, and a regulator of vascular smooth muscle cell (vSMC) differentiation^12^. The expression of SDC1 increases in endothelial cells exposed to shear stress and is differentially regulated by shear stress waveforms that mimic regions of atheroprone or atheroprotective flow in the body^13^.

While previous studies support that SDC1 can regulate cell adhesion and that the glycocalyx interacts directly with the forces of the blood flow, there is little information available that elucidates how the mechanical environment of the cells applies forces to SDC1 or regulates its binding to proteins involved in known mechanosensing pathways. In this work, we created a set of tension sensing constructs for SDC1 using a FRET-based tension sensing module consisting of Venus, mTFP1 and an elastic linker derived from the flagelliform silk protein^14,15^. Using these constructs, we examined the effects of the cellular mechanical environment on the biophysical tension in SDC1. We found that the mechanical tension in SDC1 is regulated by both substrate stiffness and the presence of nanotopographical features. In addition, changes in substrate stiffness dramatically regulate the association of SDC1 to the actin cytoskeleton, myosin IIb, Src, protein kinase A (PKA) and focal adhesion kinase (FAK). Further, the application of shear stress rapidly leads to the development of a gradient of tension in SDC1 in endothelial cells, with SDC1 under higher tension at the upstream region of the cell and under lower tension/compressive forces in the downstream region of the cell. The application of shear stress to endothelial cells rapidly altered the molecular association of SDC1 to β-actin, myosin IIb and Src. Together, our findings support that SDC1 directly responds to forces in the cellular microenvironment, leading to alterations in its binding to actin and focal adhesion-related signaling pathways.

## Results

### Validation of SDC1 FRET Tension Sensor

We created a SDC1 tension sensing construct including a full length SDC1 with the tension sensing module inserted into the ectodomain (SDC1TS). In addition, we created SDC1 tension sensing constructs with an ectodomain deletion (ΔEcto), deletion of multiple glycosylation sites (ΔGAG) and a cytoplasmic domain deletion (ΔCyto; **Fig. 1A**). Stable endothelial cell lines expressing each of the constructs were created using lentiviral transduction and puromycin selection. Cells from each line were lysed and western blotting confirmed that the molecular weight of the expressed proteins matched the expectation for the constructs and mutated proteins (**Supplemental Fig. 1**). The cellular localization of the full length construct was similar to that of endogenous SDC1, supporting appropriate membrane insertion and trafficking of the modified protein occurred (**Supplemental Fig. 2**). The ΔEcto and ΔGAG were also expressed robustly in the cells and appear to have significant membrane localization. The ΔCyto construct exhibited decreased expression and lower FRET in comparison to the other constructs, suggesting cytoplasmic domain mutant tension may possess a lower efficiency of folding or enhanced shedding. On glass culture surfaces, there was a significantly higher tension (lower FRET index) in the full length SDC1TS in comparison to the ΔEcto and ΔGAG constructs (**Fig. 1B-D**).

**Figure 1.**
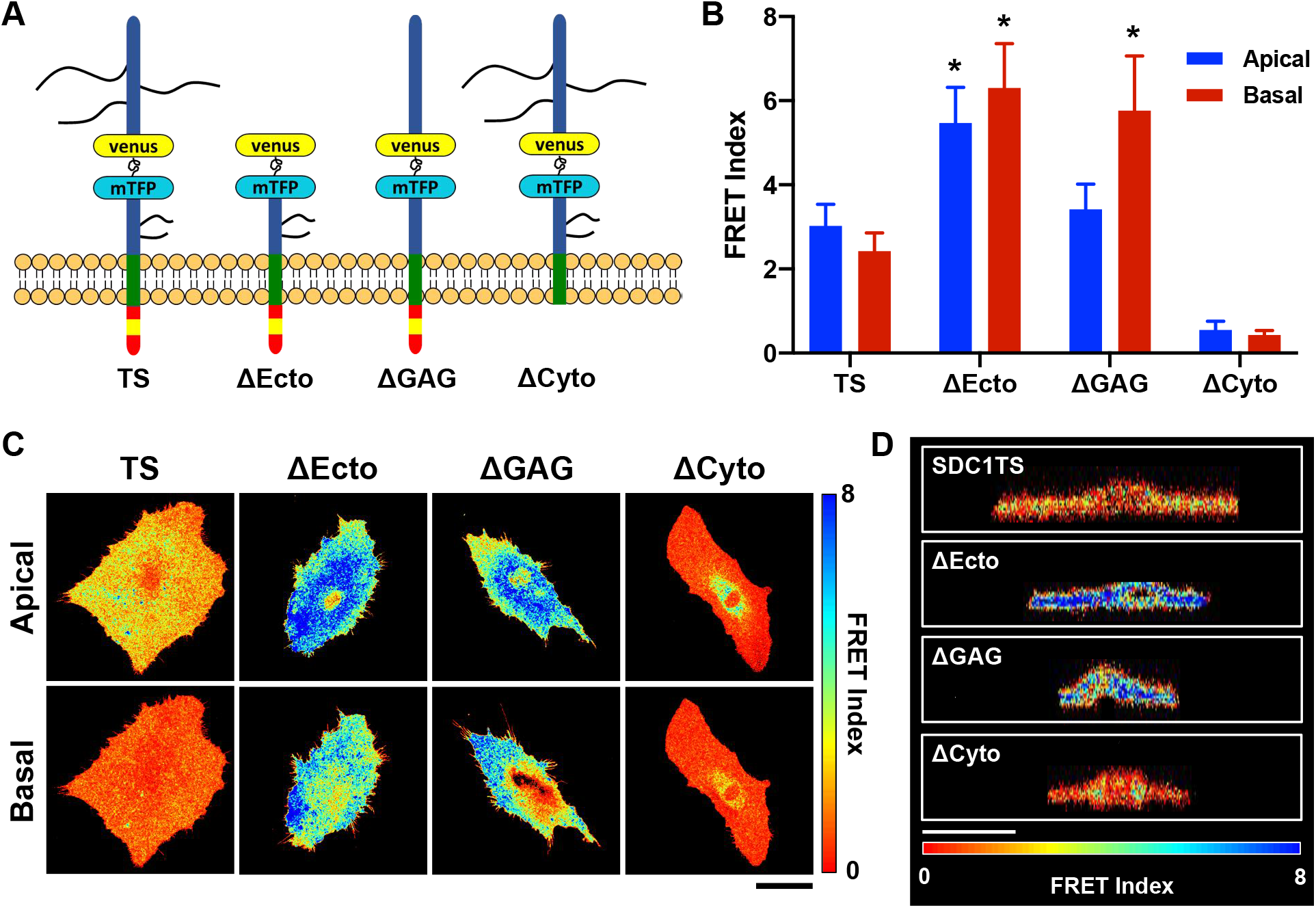
Validation of syndecan-1 FRET tension sensing constructs. (A) Diagram of SDC1 tension tensing constructs including the full length SDC1 construct (SDC1TS), ectodomain deletion mutant construct (ΔEcto), construct with mutations to remove glycosylation site (ΔGAG), cytoplasmic domain deletion mutant (ΔCyto). (B) Baseline FRET index measurements for endothelial cells grown on glass substrates. \**p* < 0.05 versus SDC1TS group. (C) FRET index images for endothelial cells expressing the different constructs grown on glass substrates. Top view of apical level (surface) and basal level (substrate) were shown. Scale bar = 50 μm. (D) Orthogonal view of reconstructed z-stacks for endothelial cells expressing the constructs. Scale bar = 50 μm.

### Tension in SDC1 is modulated by micropatterned surfaces and nanotopographical cues

We next examined whether the tension in SDC1 could be regulated extracellular matrix with different topographical cues. We cultured cells expressing the different constructs on surfaces micropatterned with crossbow and Y-shaped adhesion patterns of three different sizes **(Fig. 2A)**. The FRET index was measured on linescans taken through middle line of cells grown on the medium crossbow pattern. We observed an increase in tension of SDC1 (reduction in FRET index) in the regions of the cells that were adherent (pink areas) in comparison to non-adherent regions **(Fig. 2B)**. This pattern was not present in the other constructs that did not have the full length, glycosylated SDC1 tension sensor. We next cultured the cell lines on nanopatterned surfaces that had alternating nanogroove structures that were 800 nm in width and 500 nm in depth (**Fig. 2C**). Increased tension of SDC1 was observed in cells cultured on the nanopatterned surfaces in comparison to flat surfaces for the SDC1TS, ΔEcto and ΔCyto cell lines (**Fig. 2D)**. For the SDC1TS, ΔEcto and ΔGAG tension-sensing constructs, we found that the cells had alternating regions of high/low FRET corresponding to the spacing of nanogrooves (**Fig. 2E-G**).

**Figure 2.**
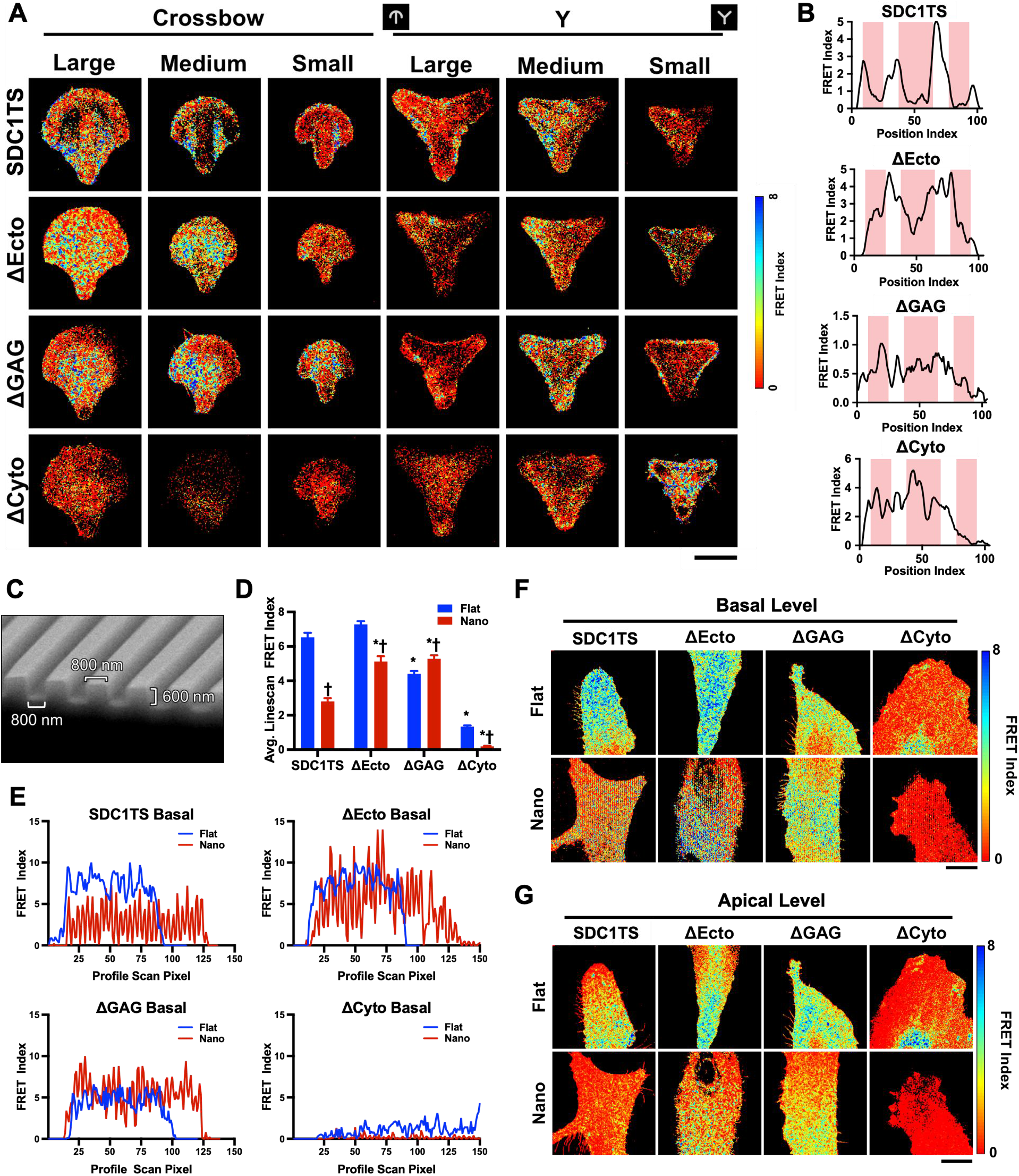
Tension in SDC1 is increased in areas of adhesion and modulated by nanotopographical cues. (A) FRET index images for endothelial cells grown on micropatterned surfaces. Scale bar = 25 μm. (B) Line scan across the middle of cells grown on the medium crossbow pattern. Pink areas are regions with collagen coating. (C) Scanning electron microscopy images of nanopatterned surface. (D) Normalized average FRET index through FRET profile scan of ECs grown on flat and nanopatterned substrates at basal level. \**p* < 0.05 versus SDC1TS on same substrate. ^†^*p* < 0.05 versus same sensor construct on flat substrate (*n* = 30 points). (E) Representative FRET index line scans perpendicular to the nanopatterned grooves at the cellular basal level. (F) FRET index images for cells at the basal level from top view. Scale bar = 50 μm. (G) FRET index images for cells at the apical level from top view. Scale bar = 50 μm.

### Substrate compliance regulates SDC1 tension and binding to cytoskeletal and specific focal adhesion signaling associated proteins

Previous studies have demonstrated that stiff substrates reduce the tight junctions and enhance permeability of endothelial monolayers through mechanism involving Src, VE-cadherin and focal adhesions^16–18^. In addition, substrate stiffness regulates endothelial inflammation and leukocyte adhesion/transmigration^16–19^. Endothelial cells also alter their mechanical stiffness based on the substrate stiffness through a mechanism involving PKA and PECAM-1^20^. To examine whether SDC1 directly responds to substrate stiffness, we cultured the endothelial cells on substrates of varying stiffness and measured the FRET within the SDC1 tension sensing constructs. We found that there was significantly decreased tension in the ΔEcto mutant tension sensor compared to the full length SDC1TS construct on soft substrates (0.2 kPa) but no significant changes were observed between other mutants and the full length tension sensor (**Fig. 3A-C**). In addition, full length SDC1TS was under higher tension in cells grown on soft substrates compared to the stiff substrates. We repeated the studies and used FRET-FLIM to confirm the changes in tension on the different substrates. The ΔEcto mutant showed decreased tension (reduction in donor lifetime) compared to full length SDC1TS, while ΔGAG mutant showed increased tension (increase in donor lifetime) compared to full length SDC1TS using FRET-FLIM imaging **(Supplemental Fig. 3)**. To examine the effect of substrate stiffness on the association of SDC1 with adhesion-related proteins, we performed pull-down assays on endothelial cells expressing HA-tagged SDC1 and grown on soft or stiff substrates. Among the proteins tested, there was a significant increase in binding to actin and myosin IIb, while there were decreases in SDC1 binding to Src, PKA and FAK in cells grown on stiff substrates versus soft substrates (**Fig. 3D, E; Supplemental Fig. 4**). Immunostaining for SDC1/Src, SDC1/β-actin and SDC1/PKA indicated that their colocalization was similarly altered by substrate stiffness (**Fig. 3F, G**).

**Figure 3.**
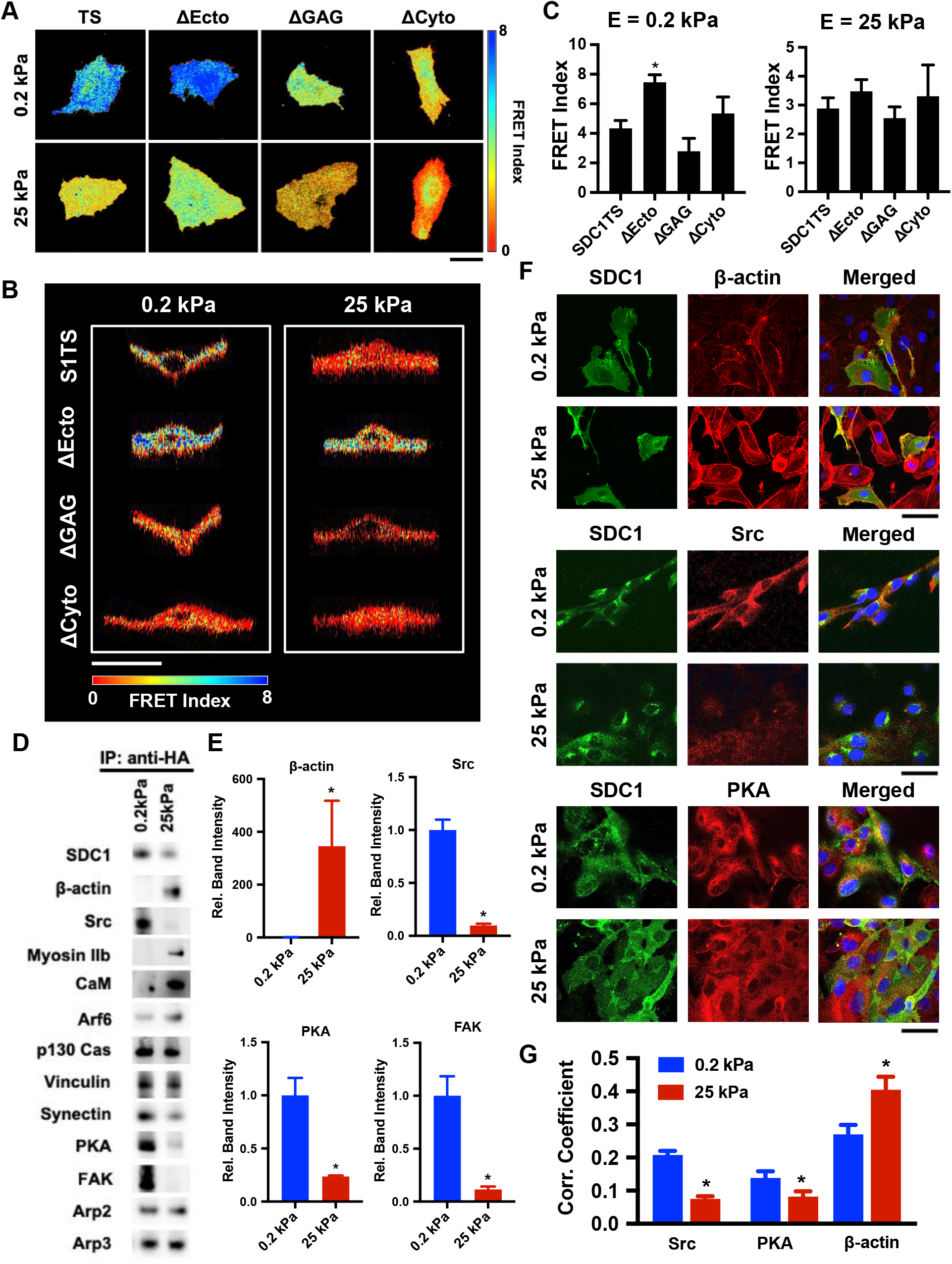
Substrate compliance regulates tension in syndecan-1 and alters its interactions with actin, Src and FAK/PKA. (A) FRET index images of cells grown on 0.2 kPa and 25 kPa substrates from top view. Scale bar = 50 μm. (B) Orthogonal view of reconstructed z-stacks for cells grown on the compliance substrates. (C) Quantification of the FRET index of cells grown on the substrates of each compliance on the basal level (*n* = 9-13). \**p* < 0.05 versus the SDC1TS on same substrate. (D) Immunoprecipitation for HA-SDC1 followed by western blotting for proteins that bind SDC1. (E) Quantification of pull-down assay relative band intensity for β-actin, Src, PKA and FAK. \**p* < 0.05 versus cells cultured on 0.2 kPa substrate (*n* = 3-6). (F) Immunostaining for SDC1/Src, SDC1/PKA and SDC1/β-actin following 24 hours of growth on the substrates of soft or stiff substrates. Scale bar = 50 μm. (I) Quantification of Pearson’s correlation coefficient of colocalization between SDC1 with Src, PKA or β-actin. \**p* < 0.05 versus cells cultured on 0.2 kPa substrate (*n* = 20-30).

### Shear stress creates a gradient in SDC1 tension and drives the association of SDC1 with actin cytoskeleton, calmodulin and Src

Endothelial cells are exposed to fluidic shear stresses from blood flow and these forces are important regulators of vascular homeostasis and arterial inflammation^21^. We next examined whether fluidic shear stress directly alters the molecular tension of SDC1 in endothelial cells. We treated the tension sensor expressing cell lines with shear stress and examined the tension in SDC1 using FRET imaging. After 15 minutes of shear stress, we found that a gradient developed in the tension of SDC1 in the cells expressing the full length construct (**Fig. 4A, B**). Along with the direction of flow, the scaled FRET index was measured from the upstream region to the downstream region of the cells (**Fig. 4C**). There was higher tension (decreased FRET index) of SDC1 in the upstream region in comparison to the downstream region. Although this observation was present in all the groups, the ratio of Front:Back FRET indexes of the three mutant groups was significantly higher than in the SDC1TS group (**Fig. 4D**). We confirmed our observation in FLIM imaging using the same shear stress setup and found consistent trends in SDC1TS and ΔCyto mutant (**Supplemental Fig. 5**). This result implies that the full length structure of SDC1 is necessary for ECs to redistribute tension under shear stress.

**Figure 4.**
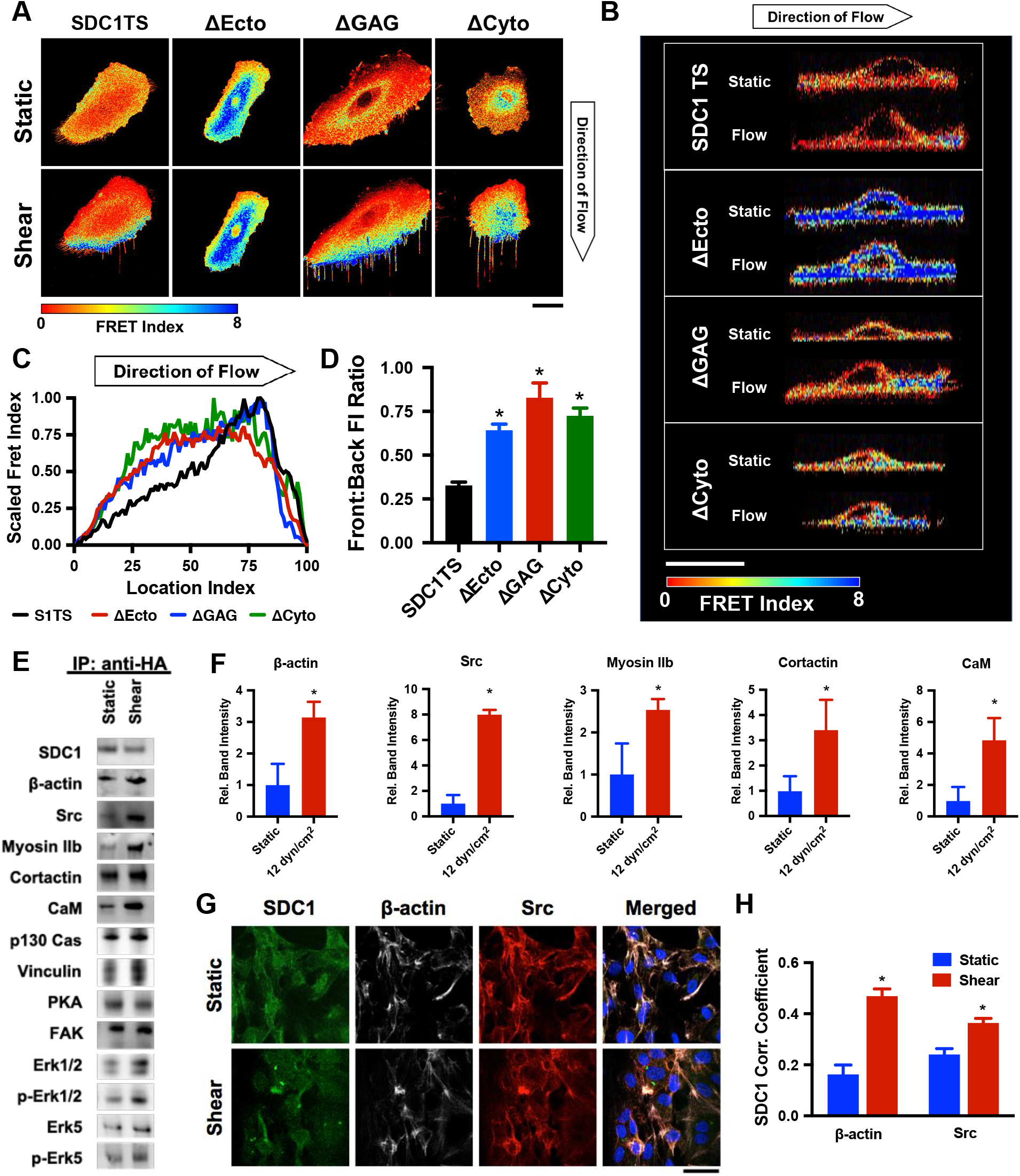
Shear stress creates a gradient in tension for SDC1. (A) FRET index images of cells following 15 minutes of shear stress at 12 dyn/cm^2^. Scale bar = 50 μm. (B) Orthogonal view of reconstructed z-stacks for the cells after 15 minutes of flow. Scale bar = 50 μm. (C) Normalized FRET index distribution across the cells following shear stress treatment (the line is an average of *n* = 5-6 cells). (D) Quantification of the FRET index ratio of the front region and back region of the cell. \**p* < 0.05 versus SDC1TS group (*n* = 5-6). (E) Immunoprecipitation of HA tagged SDC1 followed by western blotting for proteins that bind SDC1. (F) Quantification of SDC1 binding to actin, Src, myosin IIb, cortactin and CaM. \**p* < 0.05 versus static group (*n* = 3). (G) Immunostaining for SDC1, actin and Src following 15 minutes of flow at 12 dyn/cm^2^. (H) Quantification of Pearson’s correlation coefficient from colocalization analysis of SDC1 with actin and Src. \**p* < 0.05 versus cells cultured on static condition (*n* = 20-30).

We next applied 12 dyn/cm^2^ of shear stress to cells expressing HA-tagged SDC1 for 15 min and then performed immunoprecipitation on HA followed by western blotting **(Fig. 4E).** Among these proteins, we found that shear stress induced SDC1 association with β-actin, Src, non-muscle myosin IIb, cortactin, calmodulin (CaM) and integrin β3 (**Fig. 4F**; **Supplemental Fig. 6**). We performed immunostaining and found that the colocalization of Src and β-actin with SDC1 increased in response to shear stress (**Fig. 4G-H**). In addition, there was increased colocalization between SDC1 and integrin β3 but not integrin β5 (**Supplemental Fig. 7**). To investigate if activation of Src was necessary for flow-induced binding between Src and SDC1, we treated ECs with the Src inhibitor PP2 or a non-active control molecule (PP3) before applying shear stress and then repeated the experiment. Inhibition of Src blocked the flow-induced increase in Src binding to actin (**Supplemental Fig. 8**).

## Discussion

The interactions of the glycocalyx with the extracellular matrix and fluidic forces have been known for many years^22,23^. While force transmission through the glycocalyx is assumed to occur, there was a lack of direct evidence as to how and when components of the glycocalyx are under tension in cells. Syndecan-1, in particular, serves a multifaceted role in controlling cellular adhesion^24^, migration^25^ and the response to mechanical forces^11,12,26,27^. As SDC1 is expressed both on the apical and basal surfaces of endothelial cells, it has the potential to sense both the forces applied by flow and the microenvironmental cues present in the underlying extracellular matrix. In this study, we have investigated the mechanical tension and molecular interactions of SDC1 in response to extracellular matrix-mediated mechanical cues and applied fluid shear stress. Our studies support that the mechanical tension across SDC1 is responsive to changes in substrate stiffness, nanotopographical cues and to fluidic shear stresses. Moreover, our work supports that these mechanical stimuli cause alterations in the association of SDC1 with cytoskeletal and focal adhesion-related signaling pathways.

The predominant theory of the mechanisms used by cells to probe substrate rigidity consists of cells using the actomyosin cytoskeleton to deform the surrounding substrate^28^. The forces generated in this process are dependent on the substrate stiffness, leading to the generation of lower or higher forces depending on the mechanical properties of the substrate. These higher forces can lead to enhanced binding through integrin “catch bonds” and the reinforcement of the actin cytoskeleton in response to retrograde forces^29^. Syndecan-1 interacts with integrins and facilitates their binding to the extracellular matrix^24,30^. An extracellular region of SDC1 binds specifically to integrin αvβ3 and αvβ5^30^. Previous work has demonstrated that the syndecans interact with integrins in nascent focal adhesions but are not present in mature focal adhesions, supporting a model in which syndecans aid the binding of integrins but then are excluded as these integrins cluster into larger focal adhesion complexes^31,32^. Our work demonstrated greater tension in full length SDC1 in comparison to the SDC1ΔEcto mutant on soft substrates but not stiff substrates. In addition, SDC1 was not highly associated with actin or myosin IIb on the soft substrates but was associated with Src, FAK and PKA. These findings, together with previous work, suggest that, in cells on soft substrates, SDC1 binds the substrate while associated with Src/FAK/PKA and is subjected to retrograde forces from contraction of the cytoskeleton under low forces, leading to the development of tension in SDC1 and the loss of association with the actin filaments. In contrast, on stiff substrates the retraction of SDC1 occurs under higher forces leading to stabilization of its association with actin stress fibers and myosin IIb. Integrin activation in association with SDC1 leads to activation of Src, FAK and PKA, which is then dissociated from SDC1 as the integrins and Src/FAK/PKA become incorporated into a mature focal adhesions where SDC1 is excluded due to its bulky glycosaminoglycans (**Supplemental Fig. 9**).

Fluidic shear stress is known to regulate the structure of the glycocalyx^33–35^ and, conversely, specific components of the glycocalyx are required for mediating endothelial cell responses to shear stress^11,36–38^. Our group has shown in a previous study that loss of SDC1 leads to a pro-inflammatory phenotype in endothelial cells and dysfunctional mechanosensing of shear stress^11^. This study demonstrates that shear stress led to increased association of SDC1 with actin, Src, cortactin, calmodulin and non-muscle myosin IIb. Calcium signaling and Src activation are some of the earliest events in endothelial cell sensing of shear stress^39–41^. In particular, calmodulin is activated by calcium signaling and leads to phosphorylation of myosin light chain kinase (MLCK), which can subsequently regulate non-muscle myosin II. Thus, our results imply a model in which the early events of shear stress rapidly cause SDC1 to be associated with myosin IIb and actin stress fibers to provide a mechanical anchor to resist shear stress and from which tension in SDC1 can develop in the upstream edge of the cell (**Supplemental Fig. 10**). A previous study demonstrated that under shear stress the GPI-anchored protein glypican-1 redistributes downstream with shear stress whereas SDC1 does not^42^. This result is consistent with our findings that shear stress leads to dramatically increased association of SDC1 with actin and myosin IIb, effectively locking the protein in place to the cytoskeleton and allowing the tension to develop in SDC1 from the shear stress. On the downstream side of the cell there was increased FRET index in the SDC1 tension sensor, implying compressive forces on this region as the cell is sheared against substrate and the forces of flow push upon the cell membrane and underlying nucleus.

In summary, our results demonstrate that SDC1 is mechanically responsive to substrate-mediated biophysical cues and fluidic shear stresses. These findings offer a new look into the complexity of the forces being applied to the endothelial glycocalyx and delineate which molecular binding partners are modulated by substrate stiffness or shear stress among the extensive set of molecules that interact with SDC1. In response to substrate stiffness, SDC1 serves as an accessory protein facilitating the binding of integrins to the ECM, bringing Src and FAK into position for integrin-mediated signaling, and potentiating the assembly of focal adhesions in response to substrate stiffness. Under shear stress, SDC1 interacts rapidly with Src and calmodulin, suggesting its involvement in the initial steps of the response to shear stress in endothelial cells. Further studies are needed to examine the functional role of SDC1 in mechanosensing and to better understand how mechanical tension in SDC1 can alter binding to specific molecular partners.

## Acknowledgements

The authors gratefully acknowledge funding through the American Heart Association (17IRG33410888), the DOD CDMRP (W81XWH-16-1-0580; W81XWH-16-1-0582) and the National Institutes of Health (1R21EB023551-01; 1R21EB024147-01A1; 1R01HL141761-01) to ABB and R01GM112065 to JCS. The authors also acknowledge support through an American Heart Association Pre-Doctoral Fellowship to VPL (17PRE33400190).

## Materials and Methods

### Cloning of Tension Sensing Constructs

The tension-sensing module used consisted of mTFP and Venus separated by a silk flagelliform linker sequence (GPGGA)_8_, as previously described (Addgene)^43^. The open reading frame of the tension-sensing module (bases 947 to 2494) was cloned into a lentiviral expression vector containing the *Sdc-1* gene after base 965 in the *Sdc-1* gene. The ΔEcto vector was created using PCR to exclude bases 363 through 949 (amino acids 22 through 217) of the human *Sdc-1* gene. The ΔGAG vector was created using site-directed mutagenesis of serines 37, 45, 47 and 216 of the human *Sdc-1* gene into alanine residues to remove the glycosylation sites for heparan sulfate and chondroitin sulfate in the extracellular domain of *Sdc-1*^44^. The ΔCyto vector was created via site-directed mutagenesis at amino acid 278 of the human *Sdc-1* gene to create stop codons, truncating the protein to remove amino acids 279 through 310^45^. The SDC1M (mTFP; donor-only) and SDC1V (Venus; acceptor-only) constructs were created via PCR from the Sdc1TS vector template.

### Cell culture

Human umbilical vein endothelial cells (HUVECs) from a single donor were cultured in MCDB-131 culture medium with 7.5% fetal bovine serum (FBS), endothelial cell growth supplement (R&D Systems), L-glutamine and antibiotics (complete culture medium). HEK 293Ta packaging cells (ATCC) were cultured in DMEM culture medium (Gibco) supplemented with 10% heat-inactivated FBS, L-glutamine and antibiotics. Cells were incubated at 37°C with 5% CO_2_.

### Lentiviral Production and Endothelial Cell Transduction

A lentiviral packaging kit was used to produce lentiviruses (Genecopoeia). The tension sensor and controls plasmids were co-transfected with lentiviral plasmids into HEK 293Ta packaging cells according to the manufacturer’s instructions. Prior to viral transduction, HUVECs were cultured in MCDB-131 culture medium, 5% heat-inactivated FBS, L-glutamine and antibiotics. Pseudovirus-containing the lentiviral expression constructs was harvested and transduced into HUVECs in complete HUVEC culture medium containing 8 μg/ml polybrene. One day after transduction, the media was changed to complete HUVEC culture medium with 2 μg/ml puromycin. Fresh media with puromycin was added daily for 5 days to create the stable cell line.

### Immunostaining and Confocal Microscopy

Endothelial cells expressing SDC1TS, ΔEcto, ΔGAG, ΔCyto, SDC1M, SDC1V and non-transduced HUVECs at passage 8 were trypsinized and seeded at 5,000 cells/cm^2^ onto collagen-coated glass coverslips of 25 mm in diameter. Cells were cultured for 48 hours, washed with PBS at 37°C and fixed with 4% paraformaldehyde for 10 min. The cells were then washed three times with PBS and permeabilized with 0.2% Triton X-100 for 5 min. The samples were blocked with 1% BSA in PBS for 40 minutes and stained with anti-syndecan-1 (sc-12765; Santa Cruz Biotechnology) mouse and anti-GFP rabbit (ab6556; Abcam) primary antibodies diluted in PBS with 1% BSA overnight at 4°C. The cells were then rinsed with PBS with 1% BSA for 10 minutes three times. Secondary antibodies conjugated to fluorophores were added at 1:1000 dilution in 1% BSA in PBS containing 1 μg/ml of DAPI for nuclear staining. After 75 minutes of incubation at room temperature, the cells were rinsed extensively with repeated PBS washes. The coverslips were mounted onto glass slides with anti-fade mounting media (Vector Laboratories, Inc.). Fixed cells were imaged using a Fluoview FV10i confocal microscope (Olympus) at 60x magnification with oil immersion.

### Immunoblotting

Endothelial cells expressing SDC1TS, ΔEcto, ΔGAG, ΔCyto, SDC1M, SDC1V and non-transduced HUVECs at passage 8 were washed twice with cold PBS and lysed with RIPA buffer supplemented with 5 mM EDTA and protease and phosphatase inhibitor cocktail (ThermoFisher Scientific). Lysates were sonicated for 3 min, followed by centrifugation at 13,000g for 15 min. The supernatant was collected, and proteins were separated by Bis-Tris gradient (4-12%) gel (ThermoFisher Scientific) and transferred onto nitrocellulose membranes (Bio-Rad). Membranes were blocked in 5% blotto-PBST (Bio-Rad) and incubated separately with anti-syndecan-1 antibody (sc-2357; Santa Cruz Biotechnology) and anti-GFP (ab6556; Abcam) at 4C overnight. The next day, membranes were washed with PBST and incubated with HRP-conjugated species-specific antibodies (ThermoFisher Scientific) for two hours at room temperature. The stained proteins were detected by applying enhanced chemiluminescent substrate (ThermoFisher Scientific) to the membranes for 1 minute and immediately imaged using the GBox Chemi XX6 Imager.

### Spectral Imaging

Cell culture media was replaced with warm PBS, followed by HEPES-buffered Tyrode’s solution (Alfa Aesar, Inc.) prewarmed to 37°C. Spectral imaging was performed using a Zeiss 710 confocal microscope at 40x magnification with immersion oil on a stage heated to 37°C. An HFT 458/514 beam splitter was used. For spectral emission analysis, three images were obtained. For the donor (mTFP1) channel image, cells were excited with a 458 nm laser and emission was collected between 470-499 nm. For the acceptor (Venus) channel image, excitation was performed with a 514 nm laser and emission was collected between 530-600 nm. The FRET channel image was obtained using excitation of the donor and acceptor emission. Z-stacks were obtained in 0.50 μm steps.

### FRET Analysis and Image Processing

Donor, acceptor and FRET images were loaded into the FRET Analyzer and Colocalization Analyzer plugin for Fiji^46^. Bleed-through correction was performed using the Sdc1M and Sdc1V cell lines cultured on the various substrates utilized. For each slice in a z-stack, apparent FRET from non-colocalized mTFP and Venus were removed within the plugin. For cells cultured on the nanopatterned substrate, which produces autofluorescence, background-subtraction for each channel and z-position was performed before FRET-indexed images were generated. The default “Fire” lookup table (LUT) assigned by the plugin was changed to a modified “Jet” LUT. Z-stacks, z-projections, side views and 3D projections were created using Fiji software. Z-projections were generated by average intensity. Masks were created in Photoshop (Adobe) using Venus channel images to distinguish zero-FRET from regions where the sensor was not present.

### FRET Index Quantification from Images

Basal-level and apical-level slices were identified from orthogonal views of z-stacks, and average z-projections were generated. For the quantification of FRET index, cells were subdivided lengthwise, and the first and last third of the cell was traced, avoiding the nucleus. The mean FRET index was obtained from these traced regions. For cells grown on substrates of varying stiffness, FRET index was measured instead from side views (in triplicate) due to deformation of 0.2 kPa substrates by the cells. For shear stress experiments, the FRET index was scaled to the maximum value for the individual cell and then the distribution was scaled along the length axis of the cell parallel to the applied flow.

### Cell culture substrates

Unless otherwise noted, cells were seeded onto glass-bottom dishes (MatTek). The glass-bottom dishes were pre-coated with a solution of 11% (w/v) type 1 rat tail collagen (1:80 dilution, Corning) overnight at 4°C.

### Compliance substrates

For substrate compliance experiments, hydrogels (elastic moduli of 0.2 kPa and 25 kPa, Matrigen) bound to glass coverslips and pre-coated with type 1 collagen by the manufacturer were utilized. The coverslips were gently rinsed with PBS pre-warmed to 37°C and inverted onto a glass-coverslip with additional Tyrode’s solution for spectral imaging 24 to 48 hours after seeding.

### Nanopatterned substrates

For nanopatterning experiments, polyurethane nanopatterned (800:800 nm groove:gap ratio, 400 nm groove depth) and unpatterned substrates with a stiffness of 2.4 GPa (Nanosurface Biomedical, Inc.) were plasma coated to facilitate protein absorption. The substrates were then coated with a solution of 11% (w/v) type 1 rat tail collagen (1:80 dilution, Corning) overnight at 4°C. 24-48 hours after seeding, the flat and nanopatterned substrates were gently rinsed with warm PBS and Tyrode’s solution and inverted onto a glass-coverslip with additional Tyrode’s solution for spectral imaging.

### Micropatterned substrates

Micropatterned glass coverslips (CYTOOchip, Cytoo, Inc.) containing crossbows and Y shapes accommodating 700, 1100, and 1600 μm^2^ of cell area (small, medium and large, respectively) were coated with 20 μg/ml type 1 rat tail collagen according to the manufacturer’s instructions. Between 1.5 and 4 hours after seeding, micropatterned coverslips were transferred to a coverslip holder and gently rinsed with warm PBS. Spectral imaging was performed in Tyrode’s solution as described above.

### Application of shear stress

Slides containing six flow channels (μ-slide VI0.4, IBIDI, Inc.) were coated with a solution of 11% (w/v) type 1 rat tail collagen (1:80 dilution, Corning) overnight and seeded with HUVECs expressing tension sensing constructs. The cell culture medium was aspirated and replaced with Tyrode’s solution pre-warmed to 37°C. A 60 ml syringe filled with pre-warmed Tyrode’s solution was attached to silicone tubing and connected to the flow channel slides. Shear stress was applied to the channels using a syringe pump. Spectral imaging of cells under static conditions was performed with the flow system attached to the channels but without the induction of flow. Following imaging under static conditions, flow was initiated, and spectral imaging was immediately repeated under static (static-static) or flow conditions (static-12 dynes/cm^2^). For 12 dynes/cm^2^ shear stress, 9.46 ml/min flow was used, across three syringes connected to the inflow via a syringe scaffold.

### FLIM Acquisition

Cells were exposed to shear stress as described previously or seeded to compliance substrates before imaging and analyzed for fluorescence lifetime. The FLIM data were acquired and processed by the TCS SP8 STED 3X nanoscope from Leica Microsystems. Donor average lifetime of the ROIs (n ≥ 6) was recorded to compare lifetime between groups.

### Immunoprecipitation and Western Blotting

Cells were washed once with cold PBS and lysed with lysis buffer that contains 25 mM Tris, 150 mM NaCl, 1 mM EDTA, 1% Triton, 1 mM EDTA and protease and phosphatase inhibitor cocktails (ThermoFisher Scientific) for 10 min on ice. Lysates were sonicated for 3 min, followed by centrifuged at 13,000g for 10 min at 4C. The supernatant was collected and anti-HA magnetic beads (ThermoFisher Scientific) were added to the lysates followed by incubation for 30 min at room temperature on a tube rotator. Beads were collected with a magnetic stand and eluted in 1% SDS buffer. The elution samples were heated in boiling water for 10 min. Magnetic beads were removed and supernatants were saved as immunoprecipitation lysates. The proteins were separated by Bis-Tris gradient (4-12%) gel (ThermoFisher Scientific) and transferred onto nitrocellulose membranes (Bio-Rad). Membranes were blocked in 5% blotto-PBST (Bio-Rad) and incubated with primary antibodies **(Supplemental Table 1)** at 4C overnight. The next day, membranes were washed in PBST and incubated with species-specific HRP antibodies (ThermoFisher Scientific) at room temperature for two hours. The stained proteins were detected by applying ultra-sensitive enhanced chemiluminescent substrate (ThermoFisher Scientific) to the membranes for 1 minute and immediately imaged using the GBox Chemi XX6 Imager.

### Statistical Analysis

All results are shown as mean ± standard error of the mean. Comparisons between only two groups were performed using a 2-tailed Student’s t-test. Differences were considered significant at *p*<0.05. Multiple comparisons between groups were analyzed by 2-way ANOVA followed by a Tukey post-hoc test. A 2-tailed probability value <0.05 was considered statistically significant. Comparisons between only two groups, for which relative FRET is shown (normalized to baseline FRET), were performed using a Wilcoxon matched-pairs signed rank test. Differences were considered significant at *p*<0.05. Comparisons between only two unpaired groups with relative FRET values, were performed using a Mann-Whitney test. Differences were considered significant at *p*<0.05. Multiple comparisons between groups were analyzed by a Kruskal-Wallis test followed by Dunn’s post-hoc test. A 2-tailed probability value <0.05 was considered statistically significant.

## Supplemental Figure Legends

**Supplemental Figure 1.**
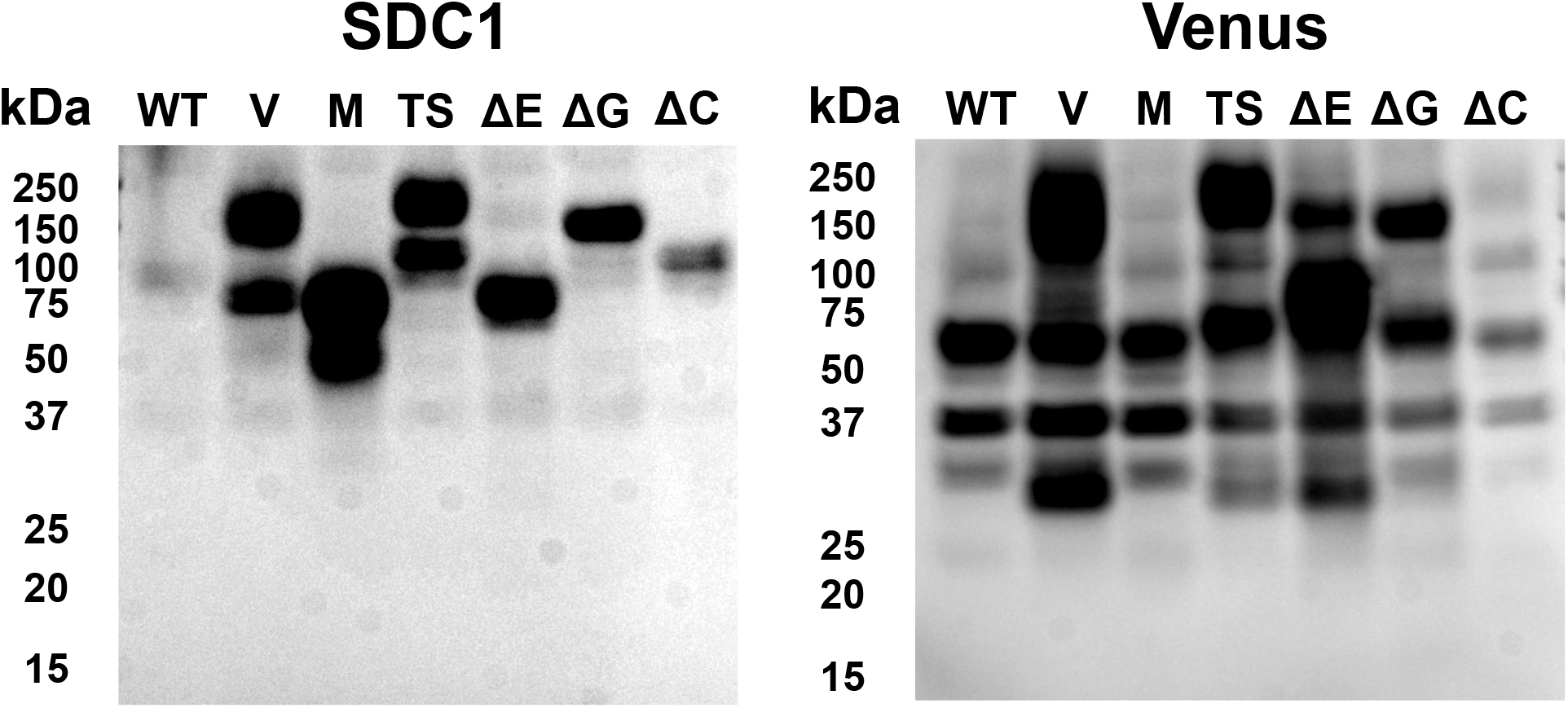
Western blotting for SDC1 and Venus in endothelial cells expressing the tension sensing constructs. Included in the blot are endothelial cells expressing the following constructs: non-transduced control cells (WT), Venus only (V), mTFP only (M), full length SDC1 tension sensing construct (TS), SDC1 TS construct with deleted ecto domain (ΔE), SDC1 TS construct with deleted glycosylation sites (ΔGAG), SDC1 TS construct with deleted cytoplasmic domain (ΔC).

**Supplemental Figure 2.**
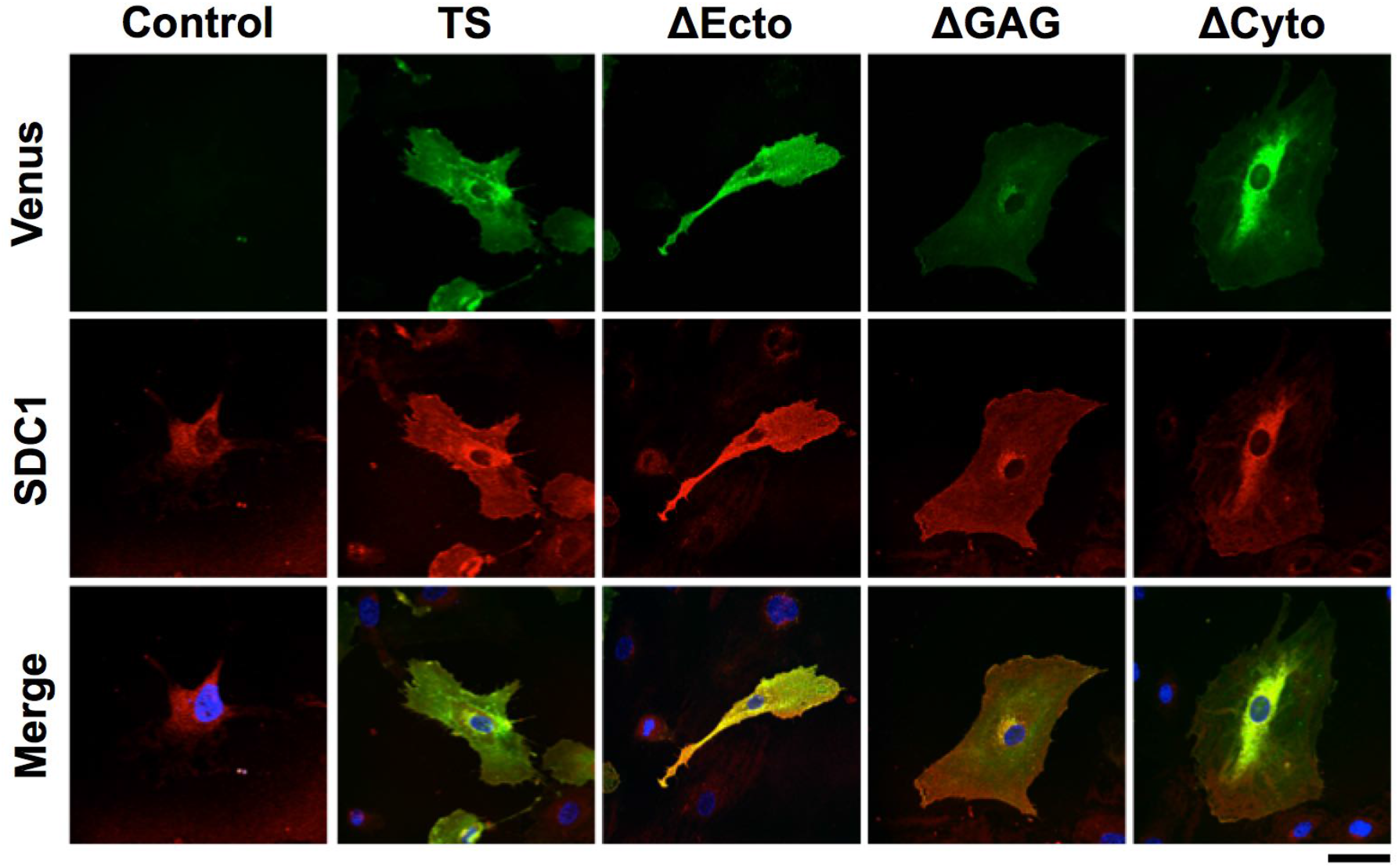
Immunostaining for Venus and SDC1 in non-transduced ECs and ECs expressing the tension sensing constructs grown on glass substrates. Scale bar = 50 μm.

**Supplemental Figure 3.**
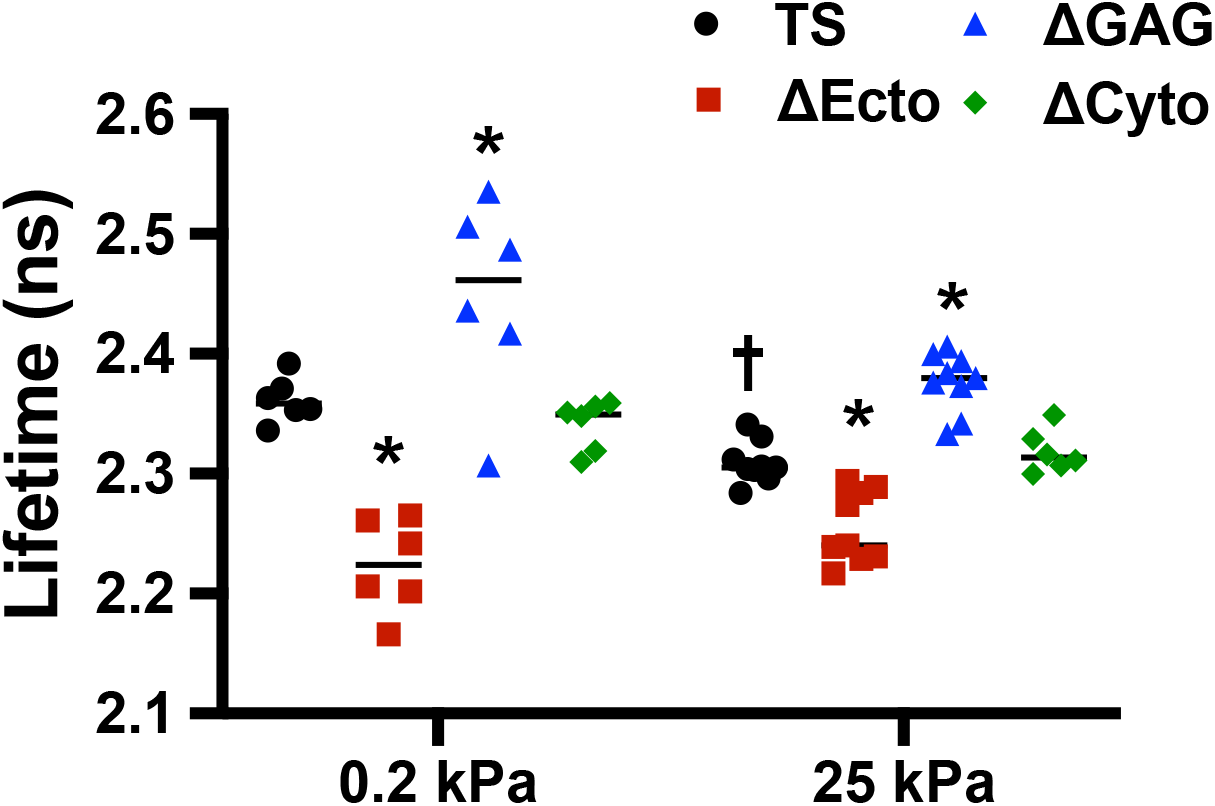
Measurements of fluorescence lifetime of donor fluorophores in cells cultured on 0.2 kPa and 25 kPa substrates. \**p* < 0.05 versus SDC1TS cultured on the substrate with same stiffness (*n*=5-9). ^†^*p* < 0.05 versus SDC1TS cultured on 0.2 kPa substrate (*n*=5-9).

**Supplemental Figure 4.**
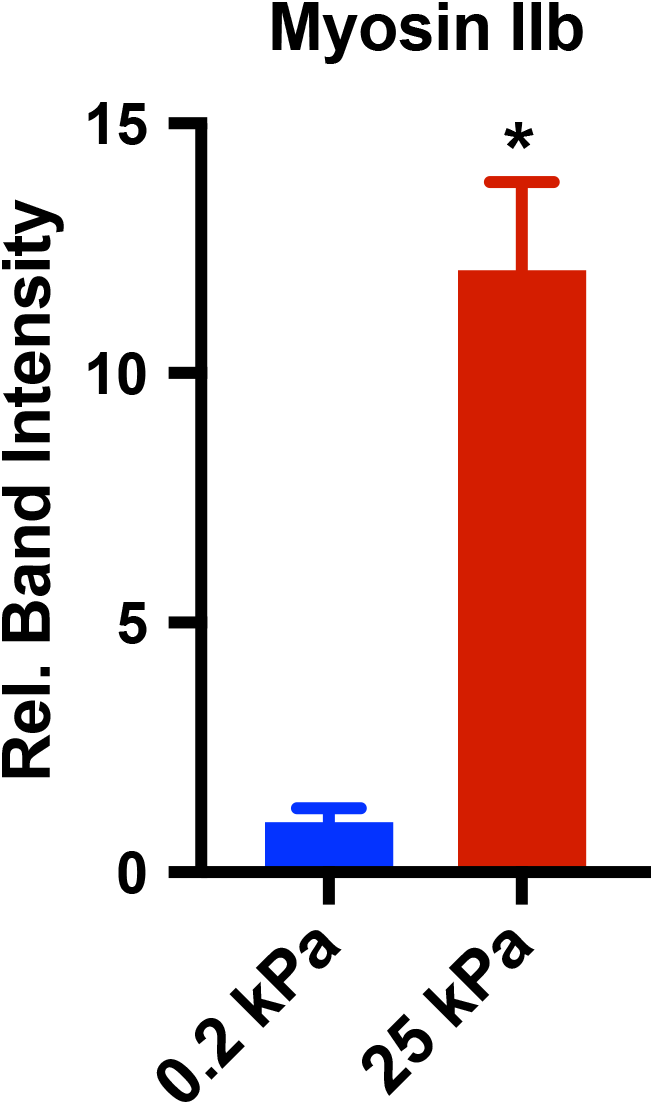
Quantification of relative band intensity on lysates from endothelial cells expressing HA-SDC1 following immunoprecipitation for HA and western blotting to myosin IIb.

**Supplemental Figure 5.**
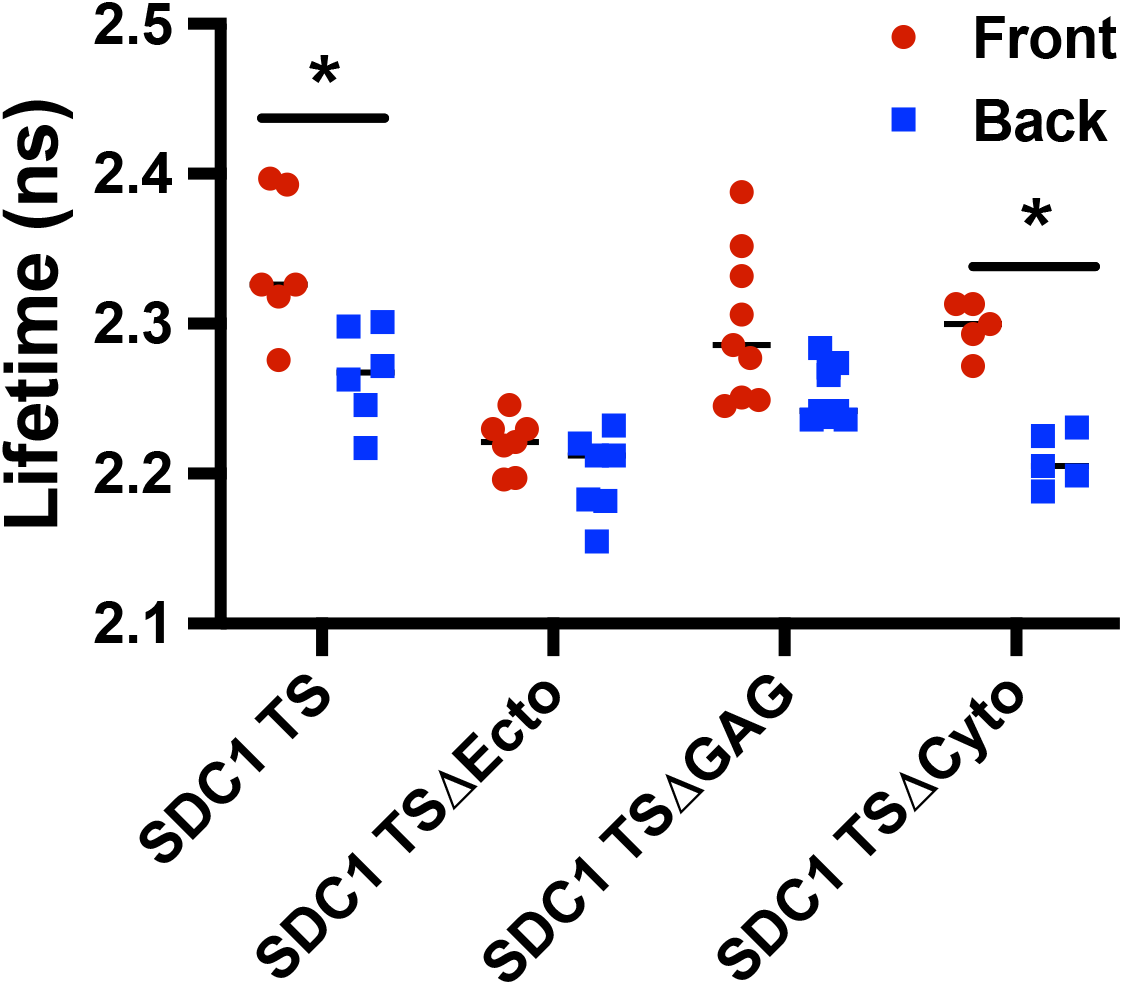
Measurements of fluorescence lifetime of donor fluorophores in the upstream region (front) and downstream region (back) of cells cultured under 12 dyn/cm^2^ of shear stress. \**p* < 0.05 versus the downstream region within the same cell line (*n*=5-9).

**Supplemental Figure 6.**
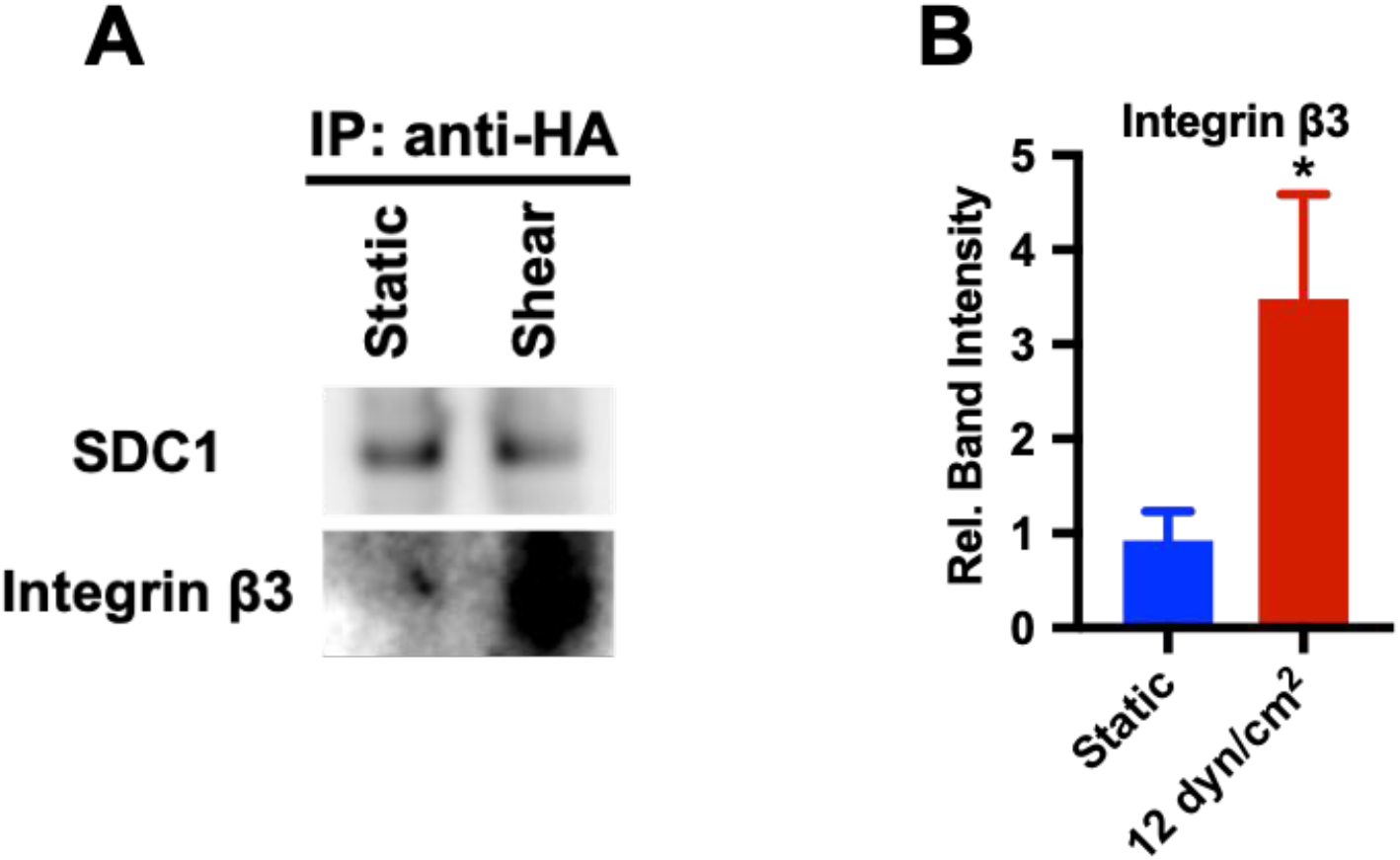
(A) Immunoprecipitation of HA tagged SDC1 in ECs after 12 dyn/cm^2^ of shear stress for 15min, followed by western blotting for integrin β3. (F) Quantification of SDC1 binding to integrin β3. \**p* < 0.05 versus cells cultured on static condition (*n* = 3).

**Supplemental Figure 7.**
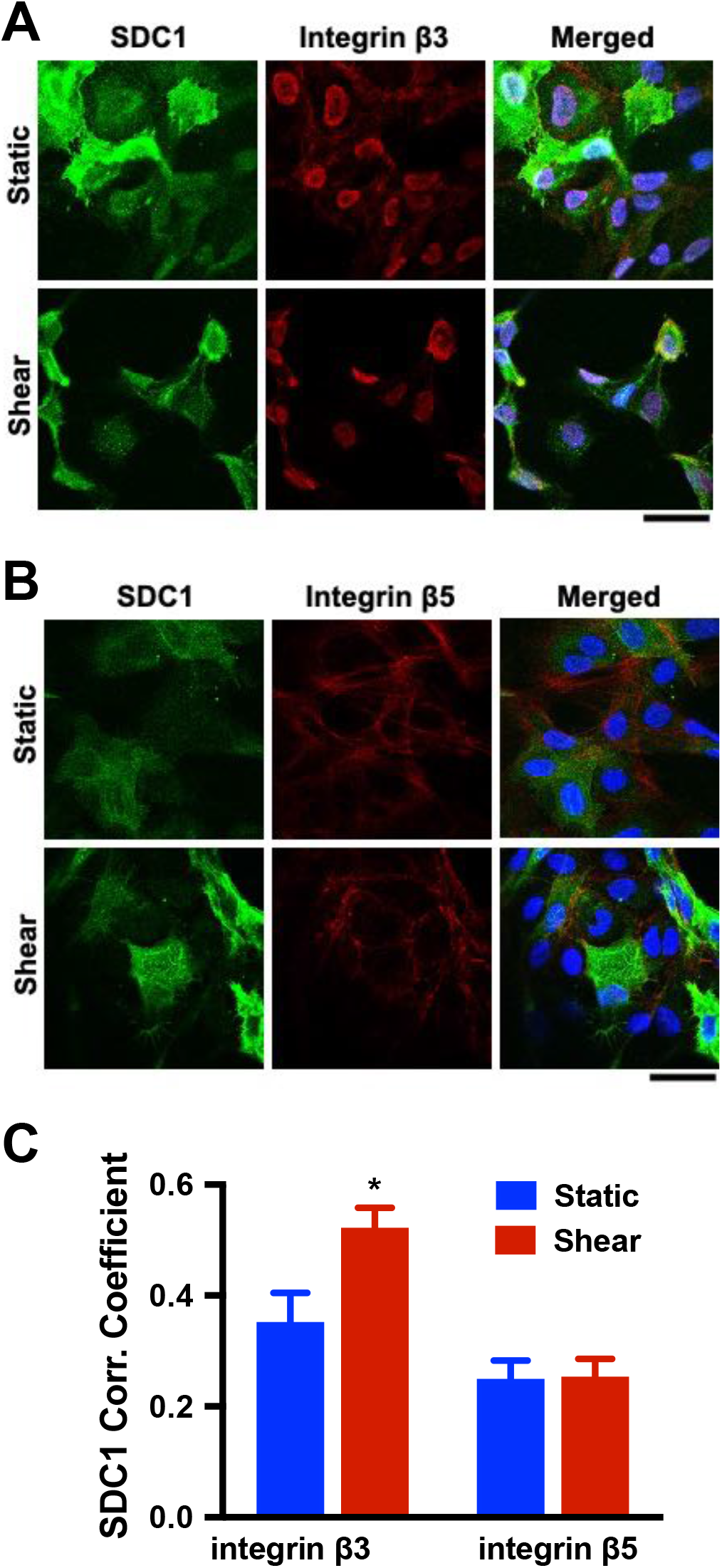
(A-B) Immunostaining for SDC1, integrin β3 and integrin β5 following 12 dyn/cm^2^ of shear stress for 15 minutes. (C) Quantification of Pearson’s coefficient in colocalization analysis of SDC1 with integrin β3 and integrin β5. \**p* < 0.05 versus cells cultured on static condition (*n* = 20-30).

**Supplemental Figure 8.**
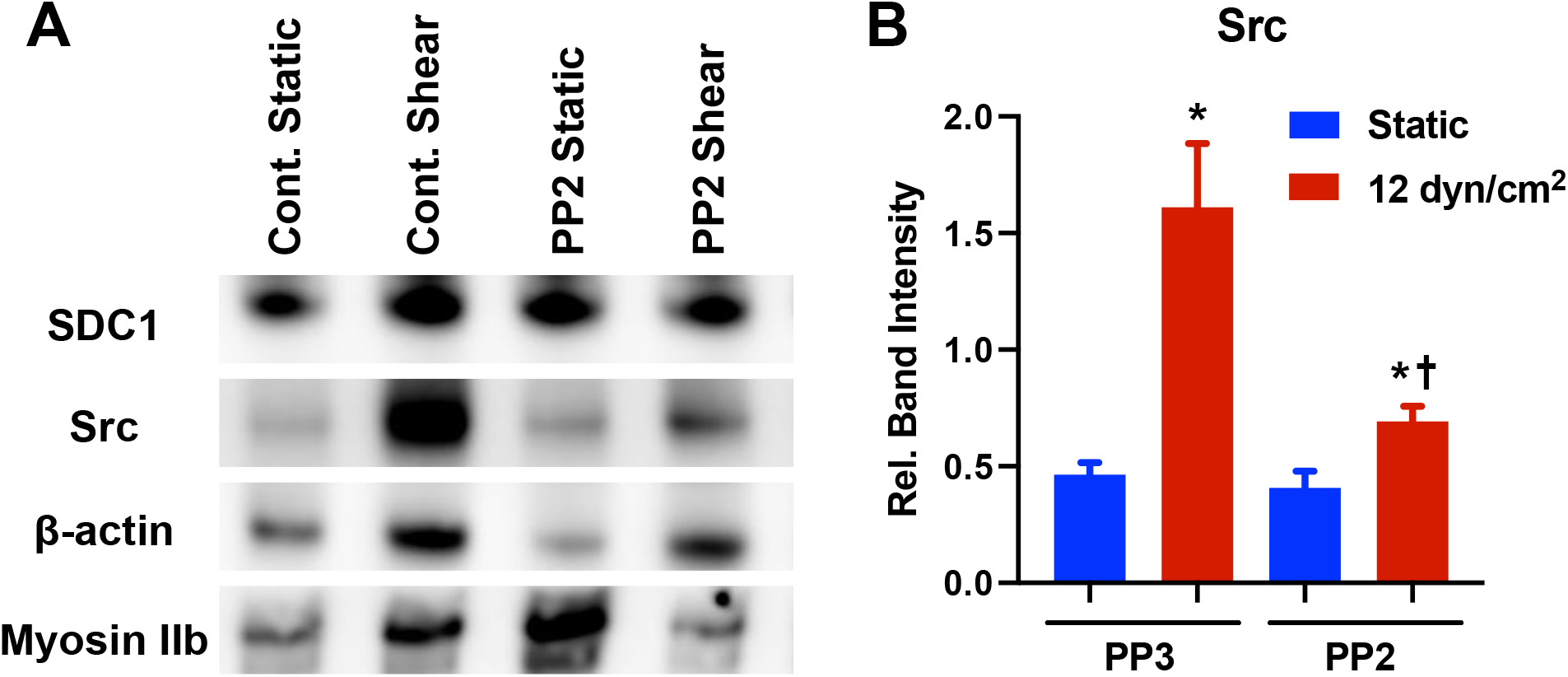
Src inhibitor PP2 inhibited SDC1 binding to Src under shear stress. (A) Immunoprecipitation of HA tagged SDC1 followed by western blotting for proteins associated with Src activation. (B) Quantification of SDC1 binding to Src. \**p* < 0.05 versus cells cultured on static condition (*n* = 3).

**Supplemental Figure 9.**
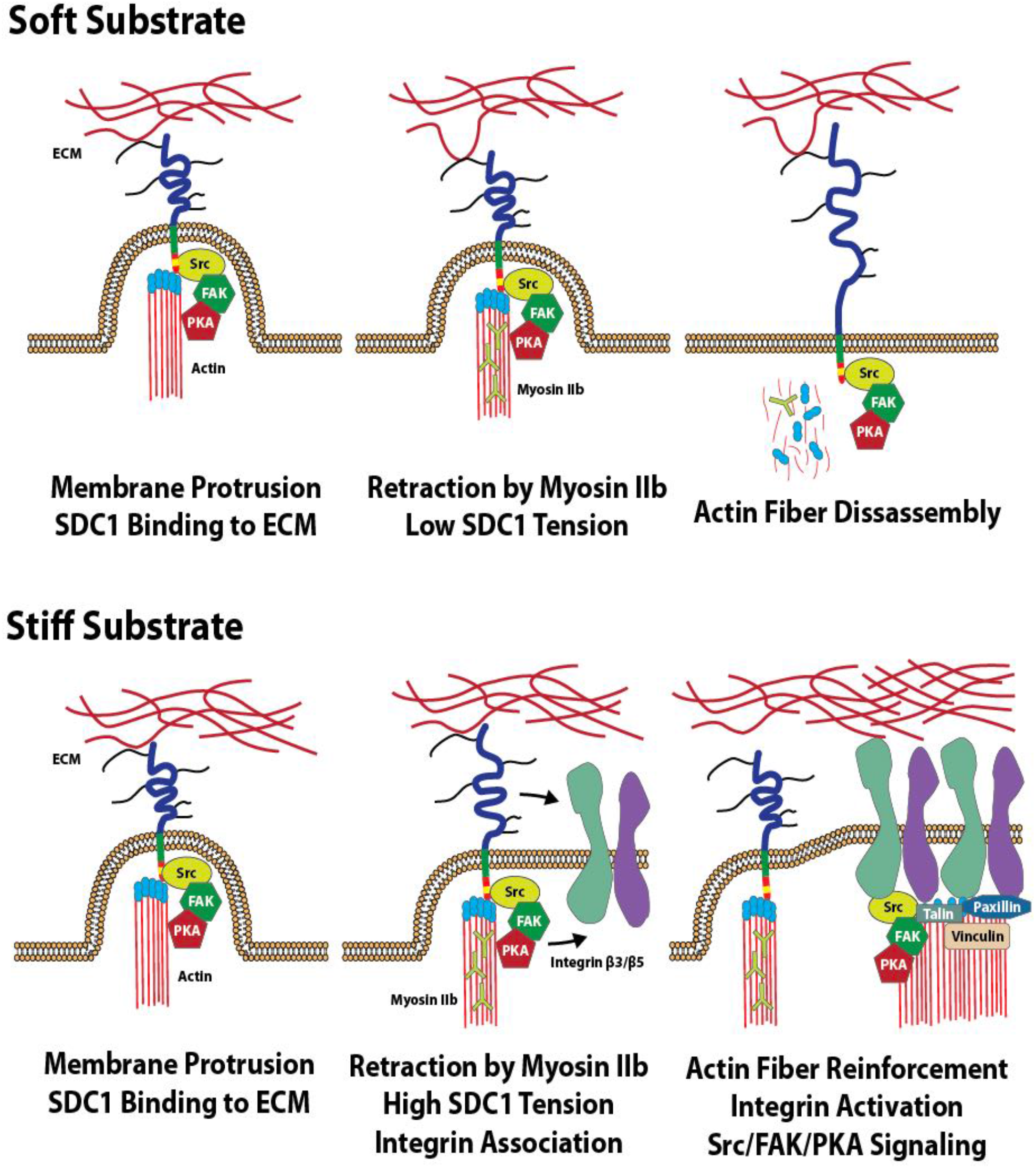
Model for involvement of SDC1 in mechanosensing of substrate stiffness in endothelial cells.

**Supplemental Figure 10.**
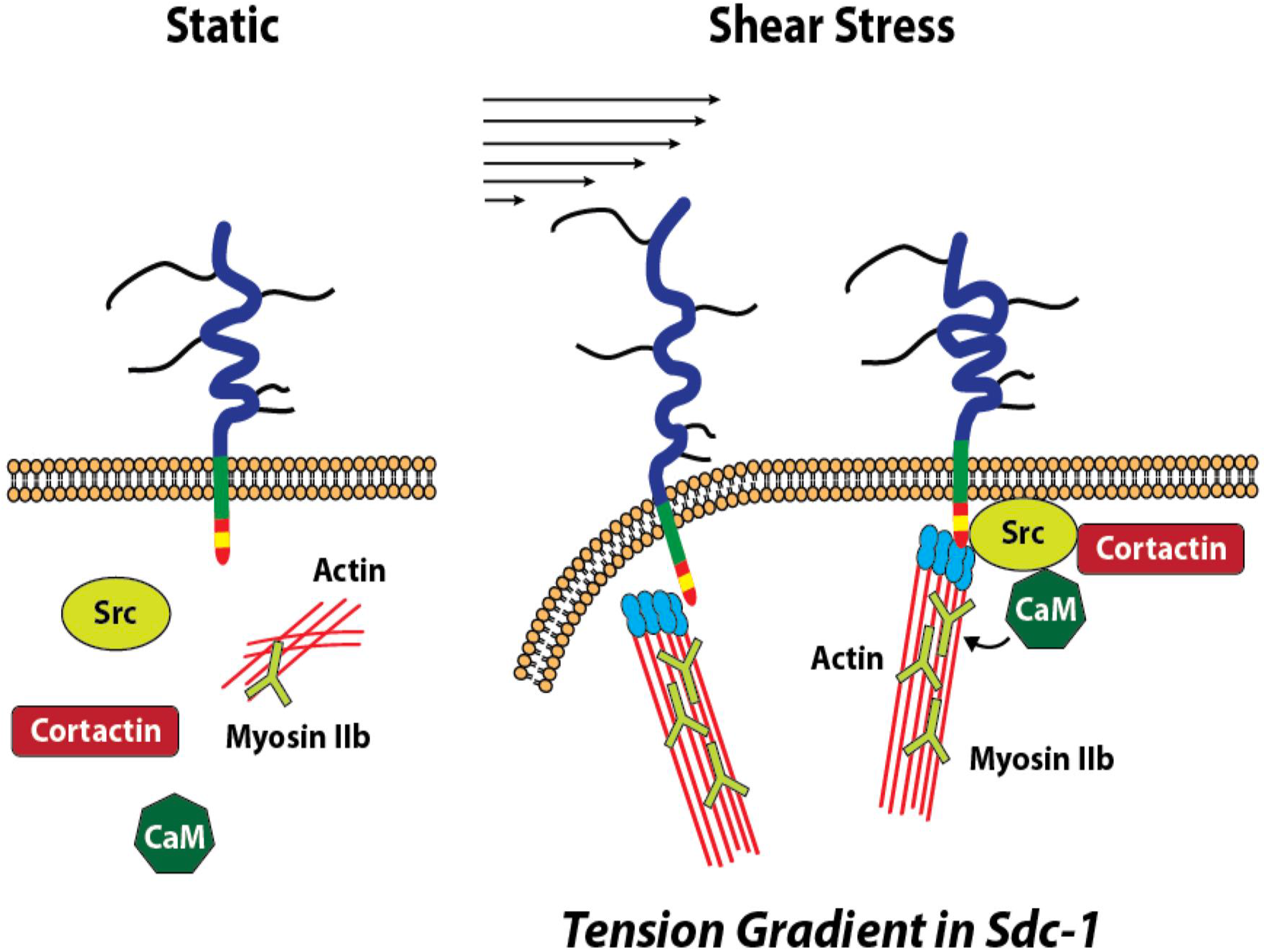
Diagram summarizing the changes in binding and tension for SDC1 under shear stress.

**Supplemental Table 1.** Primary Antibodies Used for Western Blots

| **Target Protein** | **Company** | **Catalog #** | **Species/Isotype** | **Dilution Ratio** |
| --- | --- | --- | --- | --- |
| Syndecan-1 | Cell Signaling Technology | 12922S | anti-rabbit | 1:1000 |
| β-actin | Santa Cruz Biotechnology | sc47778 | anti-mouse | 1:250 |
| Src | Cell Signaling Technology | 2108S | anti-rabbit | 1:1000 |
| Phospho-Src | Cell Signaling Technology | 2101S | anti-rabbit | 1:1000 |
| Myosin IIb | Cell Signaling Technology | 3404S | anti-rabbit | 1:1000 |
| Integrin β3 | Cell Signaling Technology | 4702S | anti-rabbit | 1:1000 |
| Integrin β5 | Cell Signaling Technology | 3629S | anti-rabbit | 1:1000 |
| Calmodulin | Cell Signaling Technology | 4830S | anti-rabbit | 1:1000 |
| p130Cas | Santa Cruz Biotechnology | sc860 | anti-rabbit | 1:250 |
| Vinculin | Cell Signaling Technology | 13901S | anti-rabbit | 1:1000 |
| RhoA | Cell Signaling Technology | 2117S | anti-rabbit | 1:1000 |
| PKA | Cell Signaling Technology | 3927S | anti-rabbit | 1:1000 |
| FAK | Santa Cruz Biotechnology | sc557 | anti-rabbit | 1:250 |
| Erk1/2 | Cell Signaling Technology | 9102S | anti-rabbit | 1:1000 |
| Phospho-Erk1/2 | Cell Signaling Technology | 9101S | anti-rabbit | 1:1000 |
| Erk5 | Cell Signaling Technology | 3372S | anti-rabbit | 1:1000 |
| Phospho-Erk5 | Cell Signaling Technology | 3371S | anti-rabbit | 1:1000 |

## References

1 Lu, D. & Kassab, G. S. Role of shear stress and stretch in vascular mechanobiology. J R Soc Interface 8, 1379–1385, doi:10.1098/rsif.2011.0177 (2011).

2 Hahn, C. & Schwartz, M. A. Mechanotransduction in vascular physiology and atherogenesis. Nat Rev Mol Cell Biol 10, 53–62, doi:10.1038/nrm2596 (2009).

3 Luo, T., Mohan, K., Iglesias, P. A. & Robinson, D. N. Molecular mechanisms of cellular mechanosensing. Nat Mater 12, 1064–1071, doi:10.1038/nmat3772 (2013).

4 Dalby, M. J., Gadegaard, N. & Oreffo, R. O. Harnessing nanotopography and integrin-matrix interactions to influence stem cell fate. Nat Mater 13, 558–569, doi:10.1038/nmat3980 (2014).

5 Discher, D. E., Janmey, P. & Wang, Y. L. Tissue cells feel and respond to the stiffness of their substrate. Science 310, 1139–1143, doi:10.1126/science.1116995 (2005).

6 Engler, A. J., Sen, S., Sweeney, H. L. & Discher, D. E. Matrix elasticity directs stem cell lineage specification. Cell 126, 677–689, doi:10.1016/j.cell.2006.06.044 (2006).

7 Kim, D. H., Provenzano, P. P., Smith, C. L. & Levchenko, A. Matrix nanotopography as a regulator of cell function. J Cell Biol 197, 351–360, doi:10.1083/jcb.201108062 (2012).

8 Swift, J. et al. Nuclear lamin-A scales with tissue stiffness and enhances matrix-directed differentiation. Science 341, 1240104, doi:10.1126/science.1240104 (2013).

9 Rahbar, E. et al. Endothelial glycocalyx shedding and vascular permeability in severely injured trauma patients. J Transl Med 13, 117, doi:10.1186/s12967-015-0481-5 (2015).

10 Sanderson, R. D., Lalor, P. & Bernfield, M. B lymphocytes express and lose syndecan at specific stages of differentiation. Cell Regul 1, 27–35, doi:10.1091/mbc.1.1.27 (1989).

11 Voyvodic, P. L. et al. Loss of syndecan-1 induces a pro-inflammatory phenotype in endothelial cells with a dysregulated response to atheroprotective flow. J Biol Chem 289, 9547–9559, doi:10.1074/jbc.M113.541573 (2014).

12 Chaterji, S., Lam, C. H., Ho, D. S., Proske, D. C. & Baker, A. B. Syndecan-1 regulates vascular smooth muscle cell phenotype. PLoS One 9, e89824, doi:10.1371/journal.pone.0089824 (2014).

13 Koo, A., Dewey Jr, C. F. & García-Cardeña, G. Hemodynamic shear stress characteristic of atherosclerosis-resistant regions promotes glycocalyx formation in cultured endothelial cells. American Journal of Physiology-Cell Physiology 304, C137–C146 (2013).

14 Grashoff, C. et al. Measuring mechanical tension across vinculin reveals regulation of focal adhesion dynamics. Nature 466, 263 (2010).

15 Conway, D. E. et al. Fluid shear stress on endothelial cells modulates mechanical tension across VE-cadherin and PECAM-1. Current Biology 23, 1024–1030 (2013).

16 Andresen Eguiluz, R. C., Kaylan, K. B., Underhill, G. H. & Leckband, D. E. Substrate stiffness and VE-cadherin mechano-transduction coordinate to regulate endothelial monolayer integrity. Biomaterials 140, 45–57, doi:10.1016/j.biomaterials.2017.06.010 (2017).

17 Kohn, J. C., Lampi, M. C. & Reinhart-King, C. A. Age-related vascular stiffening: causes and consequences. Front Genet 6, 112, doi:10.3389/fgene.2015.00112 (2015).

18 Peloquin, J., Huynh, J., Williams, R. M. & Reinhart-King, C. A. Indentation measurements of the subendothelial matrix in bovine carotid arteries. J Biomech 44, 815–821, doi:10.1016/j.jbiomech.2010.12.018 (2011).

19 Schimmel, L. et al. Stiffness-Induced Endothelial DLC-1 Expression Forces Leukocyte Spreading through Stabilization of the ICAM-1 Adhesome. Cell Rep 24, 3115–3124, doi:10.1016/j.celrep.2018.08.045 (2018).

20 Collins, C. et al. Haemodynamic and extracellular matrix cues regulate the mechanical phenotype and stiffness of aortic endothelial cells. Nat Commun 5, 3984, doi:10.1038/ncomms4984 (2014).

21 Chiu, J.-J. & Chien, S. Effects of disturbed flow on vascular endothelium: pathophysiological basis and clinical perspectives. Physiological reviews 91, 327–387 (2011).

22 Weinbaum, S., Tarbell, J. M. & Damiano, E. R. The structure and function of the endothelial glycocalyx layer. Annu. Rev. Biomed. Eng. 9, 121–167 (2007).

23 Pahakis, M. Y., Kosky, J. R., Dull, R. O. & Tarbell, J. M. The role of endothelial glycocalyx components in mechanotransduction of fluid shear stress. Biochemical and biophysical research communications 355, 228–233 (2007).

24 McQuade, K. J., Beauvais, D. M., Burbach, B. J. & Rapraeger, A. C. Syndecan-1 regulates alphavbeta5 integrin activity in B82L fibroblasts. J Cell Sci 119, 2445–2456, doi:10.1242/jcs.02970 (2006).

25 Altemeier, W. A., Schlesinger, S. Y., Buell, C. A., Parks, W. C. & Chen, P. Syndecan-1 controls cell migration by activating Rap1 to regulate focal adhesion disassembly. J Cell Sci 125, 5188–5195, doi:10.1242/jcs.109884 (2012).

26 Ebong, E. E., Lopez-Quintero, S. V., Rizzo, V., Spray, D. C. & Tarbell, J. M. Shear-induced endothelial NOS activation and remodeling via heparan sulfate, glypican-1, and syndecan-1. Integr Biol (Camb) 6, 338–347, doi:10.1039/c3ib40199e (2014).

27 Le, V. et al. Syndecan-1 in mechanosensing of nanotopological cues in engineered materials. Biomaterials 155, 13–24, doi:10.1016/j.biomaterials.2017.11.007 (2018).

28 Elosegui-Artola, A., Trepat, X. & Roca-Cusachs, P. Control of Mechanotransduction by Molecular Clutch Dynamics. Trends Cell Biol 28, 356–367, doi:10.1016/j.tcb.2018.01.008 (2018).

29 Kong, F., Garcia, A. J., Mould, A. P., Humphries, M. J. & Zhu, C. Demonstration of catch bonds between an integrin and its ligand. J Cell Biol 185, 1275–1284, doi:10.1083/jcb.200810002 (2009).

30 Beauvais, D. M., Ell, B. J., McWhorter, A. R. & Rapraeger, A. C. Syndecan-1 regulates alphavbeta3 and alphavbeta5 integrin activation during angiogenesis and is blocked by synstatin, a novel peptide inhibitor. J Exp Med 206, 691–705, doi:10.1084/jem.20081278 (2009).

31 Kechagia, J. Z., Ivaska, J. & Roca-Cusachs, P. Integrins as biomechanical sensors of the microenvironment. Nat Rev Mol Cell Biol 20, 457–473, doi:10.1038/s41580-019-0134-2 (2019).

32 Paszek, M. J. et al. The cancer glycocalyx mechanically primes integrin-mediated growth and survival. Nature 511, 319–325, doi:10.1038/nature13535 (2014).

33 Koo, A., Dewey, C. F., Jr. & Garcia-Cardena, G. Hemodynamic shear stress characteristic of atherosclerosis-resistant regions promotes glycocalyx formation in cultured endothelial cells. Am J Physiol Cell Physiol 304, C137–146, doi:10.1152/ajpcell.00187.2012 (2013).

34 Zeng, Y. & Tarbell, J. M. The adaptive remodeling of endothelial glycocalyx in response to fluid shear stress. PLoS One 9, e86249, doi:10.1371/journal.pone.0086249 (2014).

35 Li, W. & Wang, W. Structural alteration of the endothelial glycocalyx: contribution of the actin cytoskeleton. Biomech Model Mechanobiol 17, 147–158, doi:10.1007/s10237-017-0950-2 (2018).

36 Pahakis, M. Y., Kosky, J. R., Dull, R. O. & Tarbell, J. M. The role of endothelial glycocalyx components in mechanotransduction of fluid shear stress. Biochem Biophys Res Commun 355, 228–233, doi:10.1016/j.bbrc.2007.01.137 (2007).

37 Florian, J. A. et al. Heparan sulfate proteoglycan is a mechanosensor on endothelial cells. Circ Res 93, e136–142, doi:10.1161/01.RES.0000101744.47866.D5 (2003).

38 Mochizuki, S. et al. Role of hyaluronic acid glycosaminoglycans in shear-induced endothelium-derived nitric oxide release. Am J Physiol Heart Circ Physiol 285, H722–726, doi:10.1152/ajpheart.00691.2002 (2003).

39 Liu, B. et al. RhoA and membrane fluidity mediates the spatially polarized Src/FAK activation in response to shear stress. Sci Rep 4, 7008, doi:10.1038/srep07008 (2014).

40 Tzima, E. et al. A mechanosensory complex that mediates the endothelial cell response to fluid shear stress. Nature 437, 426–431, doi:10.1038/nature03952 (2005).

41 Ando, J., Komatsuda, T. & Kamiya, A. Cytoplasmic calcium response to fluid shear stress in cultured vascular endothelial cells. In Vitro Cell Dev Biol 24, 871–877, doi:10.1007/bf02623896 (1988).

42 Zeng, Y. et al. Fluid shear stress induces the clustering of heparan sulfate via mobility of glypican-1 in lipid rafts. Am J Physiol Heart Circ Physiol 305, H811–820, doi:10.1152/ajpheart.00764.2012 (2013).

43 Grashoff, C. et al. Measuring mechanical tension across vinculin reveals regulation of focal adhesion dynamics. Nature 466, 263–U143, doi:10.1038/nature09198 (2010).

44 Langford, J. K., Stanley, M. J., Cao, D. & Sanderson, R. D. Multiple heparan sulfate chains are required for optimal syndecan-1 function. J Biol Chem 273, 29965–29971, doi:10.1074/jbc.273.45.29965 (1998).

45 Miettinen, H. M., Edwards, S. N. & Jalkanen, M. Analysis of transport and targeting of syndecan-1: effect of cytoplasmic tail deletions. Mol Biol Cell 5, 1325–1339, doi:10.1091/mbc.5.12.1325 (1994).

46 Hachet-Haas, M. et al. FRET and colocalization analyzer--a method to validate measurements of sensitized emission FRET acquired by confocal microscopy and available as an ImageJ Plug-in. Microsc Res Tech 69, 941–956, doi:10.1002/jemt.20376 (2006).

